# A Murine Model for Enhancement of *Streptococcus pneumoniae* Pathogenicity Upon Viral Infection and Advanced Age

**DOI:** 10.1101/2020.07.29.227991

**Authors:** Basma H. Joma, Nalat Siwapornchai, Vijay K. Vanguri, Anishma Shrestha, Sara E. Roggensack, Bruce A. Davidson, Albert K. Tai, Anders P. Hakansson, Simin N. Meydani, John M. Leong, Elsa N. Bou Ghanem

**Affiliations:** Department of Molecular Biology and Microbiology at Tufts University School of Medicine, Boston, Massachusetts, USA; Graduate Program in Immunology, Tufts Graduate School of Biomedical Sciences, Boston, USA; UMass Memorial Health Care, University of Massachusetts Medical School, Worcester, Massachusetts, USA; Graduate Program in Molecular Microbiology, Tufts Graduate School of Biomedical Sciences, Boston, USA; Department of Anesthesiology, University at Buffalo School of Medicine, Buffalo, New York, USA; Department of Immunology, Tufts University School of Medicine, Boston, Massachusetts, USA; Department of Translational Medicine, Lund University, Malmö, Sweden; Jean Mayer USDA Human Nutrition Research Center on Aging at Tufts University, Boston, Massachusetts, USA; Stuart B. Levy Center for Integrated Management of Antimicrobial Resistance at Tufts (Levy CIMAR); Department of Microbiology and Immunology, University at Buffalo School of Medicine, Buffalo, New York, USA

**Keywords:** Co-infection, secondary bacterial pneumonia, *Streptococcus pneumoniae*, Influenza A, colonization, aging, neutrophils, inflammation

## Abstract

*Streptococcus pneumoniae* (pneumococcus) resides asymptomatically in the nasopharynx but can progress from benign colonizer to lethal pulmonary or systemic pathogen. Both viral infection and aging are risk factors for serious pneumococcal infections. Previous work established a murine model that featured the movement of pneumococcus from the nasopharynx to the lung upon nasopharyngeal inoculation with influenza A virus (IAV) but did not fully recapitulate the severe disease associated with human co-infection. We built upon this model by first establishing pneumococcal nasopharyngeal colonization, then inoculating both the nasopharynx and lungs with IAV. In young (2 months) mice, co-infection triggered bacterial dispersal from the nasopharynx into the lungs, pulmonary inflammation, disease and mortality in a fraction of mice. In old mice (20-22 months), co-infection resulted in earlier and more severe disease. Aging was not associated with greater bacterial burdens but rather with more rapid pulmonary inflammation and damage. Both aging and IAV infection led to inefficient bacterial killing by neutrophils *ex vivo*. Conversely, aging and pneumococcal colonization also blunted IFN-α production and increased pulmonary IAV burden. Thus, in this multistep model, IAV promotes pneumococcal pathogenicity by modifying bacterial behavior in the nasopharynx, diminishing neutrophil function, and enhancing bacterial growth in the lung, while pneumococci increase IAV burden likely by compromising a key antiviral response. Thus, this model provides a means to elucidate factors, such as age and co-infection, that promote the evolution of *S. pneumoniae* from asymptomatic colonizer to invasive pathogen, as well as to investigate consequences of this transition on antiviral defense.

## INTRODUCTION

*Streptococcus pneumoniae* (pneumococcus) is a Gram-positive pathobiont that typically resides asymptomatically in the nasopharynx of healthy individuals (1). It is hypothesized *S. pneumoniae* establishes an asymptomatic biofilm on the nasopharyngeal epithelium by attenuating the production of virulence factors and concomitant inflammation (2–5). However, when immunity is compromised, a common occurrence in aging (6), pneumococci can cause serious disease such as otitis media, pneumonia, meningitis and bacteremia (7). In humans, pneumococcal carriage is believed to be a prerequisite to invasive disease (8, 9). Bacterial isolates from invasive infections are genetically identical to those found in the nasopharynx of patients (9); this and other longitudinal studies have led to the suggestion that invasive disease often involves pneumococcal carriage in the upper respiratory tract (8, 9).

The rate of reported colonization is quite variable among adults and is confounded by differences in detection methods, but colonization may be more prevalent in the elderly (10). A meta-analysis of twenty nine published studies found that in individuals above 60 years of age, conventional culture showed 0-39% carriage, while 3-23% carriage was detected by molecular methods (11). Importantly, carriage was higher among nursing home residents (11). In older adults, conventional culture methods estimated carriage to be <5% (12–14) but more recent data from European studies using molecular detection methods indicated that carriage in the elderly ranges from 10-22% (10, 15–17). Thus, carriage rates may be much higher than what was previously estimated in elderly individuals (12–14).

Advanced age increases the risk of invasive pneumococcal disease and pneumococcal pneumonia (7). Aging is associated with immunosenescence, the overall decline in immunity that accompanies aging, as well as inflammaging, a low-grade chronic inflammation that render the elderly more susceptible to pulmonary infections (18). Polymorphonuclear leukocytes (PMNs), also known as neutrophils, are a crucial determinant of age-related susceptibility to primary pneumococcal infection (19). These cells are required to control bacterial burden at the start of infection (20–22). However, aging is accompanied by impaired PMN anti-bacterial function (23, 24). In addition, aged hosts experience exacerbated PMN pulmonary influx during primary pneumococcal pneumonia (19, 25, 26), and persistence of PMNs in the airways beyond the first few days leads to tissue destruction and systemic spread of infection (27, 28).

In addition to advanced age, epidemiological and experimental data show that invasive pneumococcal infections are strongly associated with viral infection (29–31). The risk of pneumococcal pneumonia is enhanced 100-fold by influenza A virus (IAV) infection (32, 33), resulting in the seasonal peak of pneumococcal disease during influenza outbreaks (33). Further, *S. pneumoniae* is historically among the most common etiologies of secondary bacterial pneumonia following influenza and associated with the most severe outcomes (30, 34–36). Symptoms of secondary bacterial pneumonia include cough, dyspnea, fever, muscle aches and when severe result in hospitalizations, respiratory failure, mechanical ventilation, and can lead to death (30, 34–36).

IAV commonly infects the upper respiratory tract, and at this location viral infection can enhance the nutritional environment for pneumococcus in the nasopharynx, leading to greater bacterial loads and/or higher rates of bacterial acquisition (4, 37). IAV can also directly bind the pneumococcal surface and enhance bacterial binding to the pulmonary epithelium leading to increased colonization (38). In addition, viral infection of the pulmonary epithelium induces the release of host components, such as adenosine triphosphate and norepinephrine, that are sensed by biofilm-associated pneumococci, triggering both the production of pneumococcal virulence factors and the dispersal of bacteria (4, 37, 39, 40). This in turn facilitates bacterial spread to and colonization of the lower respiratory tract (4).

IAV can infect not just the upper respiratory tract, but also the lungs, and murine models featuring sequential pulmonary challenge with IAV first followed by pneumococcus show the virus can compromise immune defense against pneumococcus (29, 34, 41). In these models, IAV alters both the pulmonary environment and the immune response to enhance subsequent bacterial colonization and tissue damage. For example, IAV pre-infection increases mucus production and fibrosis and dysregulates ciliary function (34, 42, 43), thus impairing mechanical clearance of invading bacteria. Viral enzymes, along with virus-elicited inflammation, result in the exposure of epithelial proteins that promote pneumococcal adherence to and invasion of host cells (41, 44–46). Further, IAV triggers type I and II interferon (IFN) responses that impair both the recruitment and antibacterial function of phagocytes key to defense against pneumococcus (47–50). The combined tissue damage and the compromise in immune function render the lung more permissive for invasive *S. pneumoniae* infection (34, 45, 47). Less understood is how *S. pneumoniae* infection may alter host antiviral responses and viral replication in the lung.

Notably, advanced age and IAV infection appear to synergistically enhance susceptibility to pneumococcal lung infections (6, 18, 51, 52). Indeed, individuals ≥ 65 years old account for 70-85% of deaths due to pneumonia and influenza (52). Interestingly, elderly individuals with influenza-like symptoms were reported to have an increased pneumococcal carriage rate of 30% (53). In animal models in which bacteria are directly instilled into the lungs following influenza infection, aging is associated with increase susceptibility of secondary pneumococcal pneumonia (51). Age-dependent changes in the expression of key components of innate immune signaling contribute to disease in this co-infection model (51). However, the factors, including those that are age-dependent, that trigger the transition of pneumococci from benign colonizer to pathogen are poorly defined, in part because small animal models that recapitulate the transition from asymptomatic colonization to overt clinical illness are lacking.

These above studies indicate that bacterial-viral synergy is multi-factorial and can occur at different sites with the host. Insight into events that occur in both the nasopharynx and lung and contribute to the heightened susceptibility of the aged to serious disease upon pneumococcal/IAV coinfection is needed to develop better therapeutic and preventative approaches. A current murine model for the spread of nasopharyngeal pneumococci to the lung after viral infection of the upper respiratory tract relies on initial bacterial colonization of the nasopharynx followed by viral infection, but does not recapitulate the severe signs of human clinical disease (4). Influenza virus is capable of infection not just of the upper respiratory tract, but also the lung (54, 55). To better investigate the transition of *S. pneumoniae* from asymptomatic colonizers to invasive pathogens following IAV infection, as well as the effect of host age on the disease process, we built upon this mouse model of *S. pneumoniae/*IAV co-infection by incorporating viral infection of the lower respiratory tract. The enhanced model recapitulates the severe and age-exacerbated clinical disease observed in humans.

## RESULTS

### Intranasal IAV inoculation of mice pre-colonized with *S. pneumoniae* strain TIGR4 does not result in disease

Biofilm-grown *S. pneumoniae* are relatively less virulent and thus adapted to host colonization rather than disease (2–4). A previously established murine model in BALB/cByJ mice utilizes biofilm-grown *S. pneumoniae* serotype 2 strain D39 and serotype 19F strain EF3030 to establish heavy carriage in the nasopharynx (4). Then, two days after bacterial inoculation, IAV is introduced into the nasal cavity and results in the spread of pneumococci from the nasopharynx (NP) to the lung (4). We first recapitulated this model with *S. pneumoniae* strain TIGR4, an invasive serotype 4 strain (56, 57) that we previously found to be highly virulent in aged C57BL/6 mice (19). We intranasally (i.n.) inoculated young (8-10 weeks) C57BL/6 (B6) mice with 1×10^6^ colony forming units (CFU) of biofilm-generated *S. pneumoniae* TIGR4 by delivering the bacteria in 10 µl (a volume that is unlikely to inoculate the lung (58)) to the nares of non-anesthetized mice. Forty-eight hours later, mice were i.n. inoculated with 10 µl containing 20 PFU of Influenza A (IAV) virus PR8. (See Fig. S1A for general scheme). Control groups of mice were either infected with *S. pneumoniae* or IAV alone as controls. However, under these conditions, mice did not display signs of sickness, nor did bacteria spread into the lungs after seven days (not shown).

To increase the likelihood of clinical disease, we repeated the experiment with a five-fold higher (5×10^6^ CFU) dose of bacteria and a 25-fold higher dose (500 PFU) of IAV. Similar to earlier reports for other *S. pneumoniae* strains (D39 and EF3030) (4), IAV co-infection resulted in a 10-fold increase in *S. pneumoniae* TIGR4 in the nasal lavage fluid at 2 days post-IAV infection (Fig. S1B). In addition, IAV co-infection was associated with the detection of bacteria in the lungs of 40% of mice, compared to none in the control group infected with *S. pneumoniae* TIGR4 alone. This trend is consistent with the previous BALB/cByJ mice model of coinfection (4), but did not reach statistical significance. Furthermore, co-infection was not associated with weight loss when assessed over the course of 4 days post-infection (Fig. S1C). We also scored mice for clinical signs of the disease based on weight loss, activity, posture and breathing and ranging from healthy [score = 0] to moribund [score = 25] and requiring euthanasia if the score was >9 as previously described (59). As secondary pneumonia can occur several days following IAV (41), we monitored the disease course up to 7 days, but did not detect disease symptoms or death in any of the co-infected mice (100% survival and 0 daily clinical score, including weight loss for all mice). Therefore, despite promoting bacterial dispersal from the nasopharynx into the lungs, similar to the previous work with other *S. pneumoniae* strains and BALB/c mice (4), this model of *S. pneumoniae* TIGR4/IAV coinfection did not result in overt clinical signs of disease.

### Combined intranasal/intratracheal IAV inoculation of *S. pneumoniae-*colonized mice results in bacterial dissemination and disease

IAV infection is not restricted to the upper respiratory tract, and can cause viral pneumonia in a significant fraction of infected individuals (54, 55) that is likely to be crucial for creating an environment in the lungs that is more permissive for bacterial infection (29, 34, 41). Indeed, viral lung infection diminishes pulmonary defenses against *S. pneumoniae* and promotes secondary bacterial pneumonia (34, 41, 44–50). Delivery of IAV i.n. to BALB/cByJ mice results in signs of viral pneumonia (4), but we found that pulmonary access of inocula delivered via the nasopharynx is more restricted in B6 mice as compared to BALB/c mice (unpublished observation), raising the possibility that the lack of disease observed in co-infected B6 mice was due to the exclusive localization of virus in the nasal cavity, with limited opportunity to alter systemic or pulmonary immunity. In fact, when we measured pulmonary viral load two days following i.n. infection with 500 PFU IAV, we were unable to detect any PFU in the lungs, while delivery of 20 PFU of IAV by intratracheal (i.t.) inoculation was sufficient to establish lung infection (Fig. S2).

Therefore, to ensure delivery of IAV to both the nasopharynx and the lungs, we co-infected *S. pneumoniae*-colonized B6 mice by delivering the virus by two routes. Mice were inoculated i.n. with 5×10^6^ CFU of biofilm grown S. *pneumoniae* TIGR4 and 48 hours later infected not only with 500 PFU IAV i.n., but also 20 PFU i.t. to ensure pulmonary infection (Fig. 1A). No mice in a control group inoculated i.n. with *S. pneumoniae* alone lost weight (Fig 1B), displayed clinical signs of sickness (Fig 1C), or died (Fig 1D). A second control group, inoculated i.n. and i.t. with IAV alone displayed no disease until after day 4, when they exhibited weight loss (Fig 1B) and began succumbing to viral infection (Fig 1D). In contrast, inoculation of IAV to animals pre-colonized with *S. pneumoniae* caused a bacterial/viral co-infection that resulted in weight loss (Fig. 1B), clinical symptoms (Fig. 1C) and death (Fig. 1D) that were detected starting day 2 post IAV introduction. At this time point, a higher fraction of co-infected mice displayed signs of disease as compared to controls infected with IAV only (55% vs 64%), and disease was more severe in mice that displayed clinical symptoms, although this did not reach statistical significance. In addition, the overall survival rate among co-infected mice was significantly lower than mice singly infected with *S. pneumoniae* alone (Fig. 1D). These signs of exacerbated disease were associated with significantly higher bacterial burdens in the nasopharynx as well as translocation of *S. pneumoniae* into the lung in comparison to mice colonized with the bacteria or mice infected with IAV alone (Fig. 1E). These findings demonstrated that the new co-infection model leads to disease in a fraction of young healthy mice, which may increase the likelihood of detecting enhanced susceptibility in vulnerable hosts.

**Figure 1.**
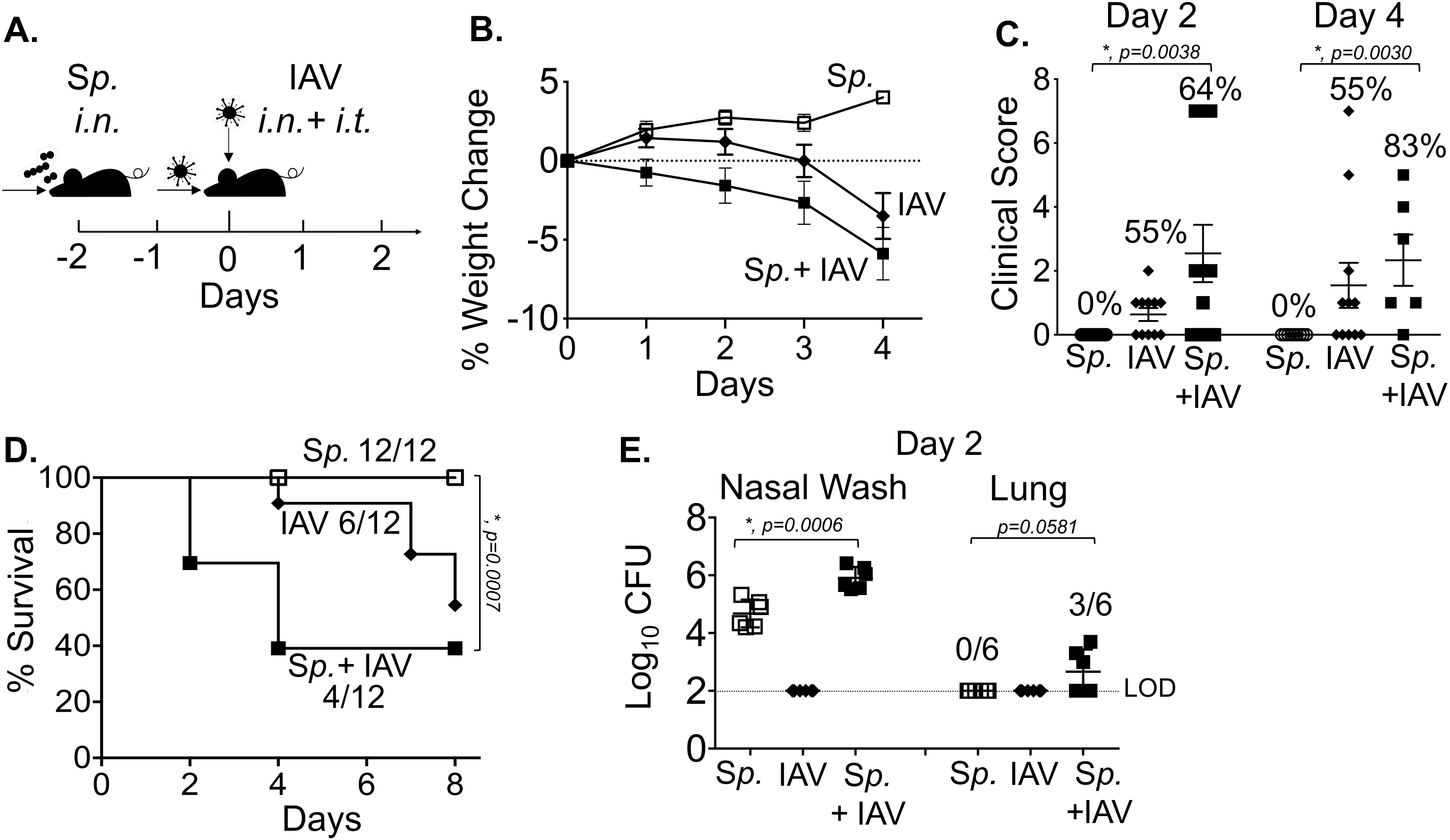
Combined intranasal/intratracheal IAV inoculation of *S. pneumoniae-* colonized mice results in bacterial dissemination and disease. (A) Timeline of co-infection; 8-10 weeks old male C57BL/6 (B6) mice were inoculated i.n. with 5×10^6^ CFU of biofilm grown S. *pneumoniae* TIGR4 to establish colonization in the nasopharynx. 48 hours later, the mice were either mock treated (*Sp*) or received 500 PFU of Influenza A virus PR8 (IAV) i.n. and 20 PFU i.t. (B) Percent weight loss was monitored daily. (C) Blinded clinical scoring was performed on day 2 and day 4 post IAV infection. The percentages denote the number of sick mice observed over total number of mice. A score of 0 means no sign of sickness observed and a score above 1 indicates observable sickness. (D) Survival was monitored for 8 days post IAV infection with fractions denoting survivors over total number of mice. (E) The bacterial burden in the nasopharynx and lung were determined at day 2 post IAV infection. Pooled data from four separate experiments are shown in which one group of twelve mice in each experimental condition were monitored over time and another group of 6 mice per experimental condition were used for measuring bacterial burden. Statistically significant differences determined by Student’s t-test for bacterial burden and clinical score and the log-rank (Mantel-Cox) test for survival are indicated by asterisks. #, indicates statistical significance (*p*< 0.05) between *Sp* and *Sp*+IAV groups by Fisher’s exact test.

### PMN depletion may have a small effect on the course of disease during IAV/*S. pneumoniae* co-infection

We previously found that in primary pneumococcal pneumonia, PMNs are required to control bacterial numbers early in the infection process; however, their persistence in the lungs is detrimental to the host and can promote the infection at later time points (20). To address the role of PMNs during co-infection, we treated young mice with PMN-depleting anti-Ly6G antibody (1A8) one day prior to pneumococcal colonization and throughout the co-infection (based on timeline in Figure 1A). We then confirmed that the cells were depleted by staining with the RB6 antibody followed by flow cytometry (Fig S3). Following infection, we measured bacterial burden, weight loss, clinical score and survival over time. PMN depletion had no effect on bacterial burdens in the nasopharynx or bacterial spread to the lungs or blood following co-infection (Fig 2A and B). However, PMN-depleted mice appeared to lose more weight at day 3 and 4 post co-infection as compared to the control group (Fig. 2C), and a greater proportion of PMN-depleted mice displayed clinical signs of sickness as compared to the control group at both 18 hours and 48 hours post IAV infection (Fig. 2D). Additionally, lower survival was observed in the PMN-depleted group, in which 25% survived to day 7 compared to ∼43% in the untreated control group (Fig. 2E). These differences did not reach statistical significance, but raise the possibility that PMNs provide a measure of defense to co-infection in young mice.

**Figure 2.**
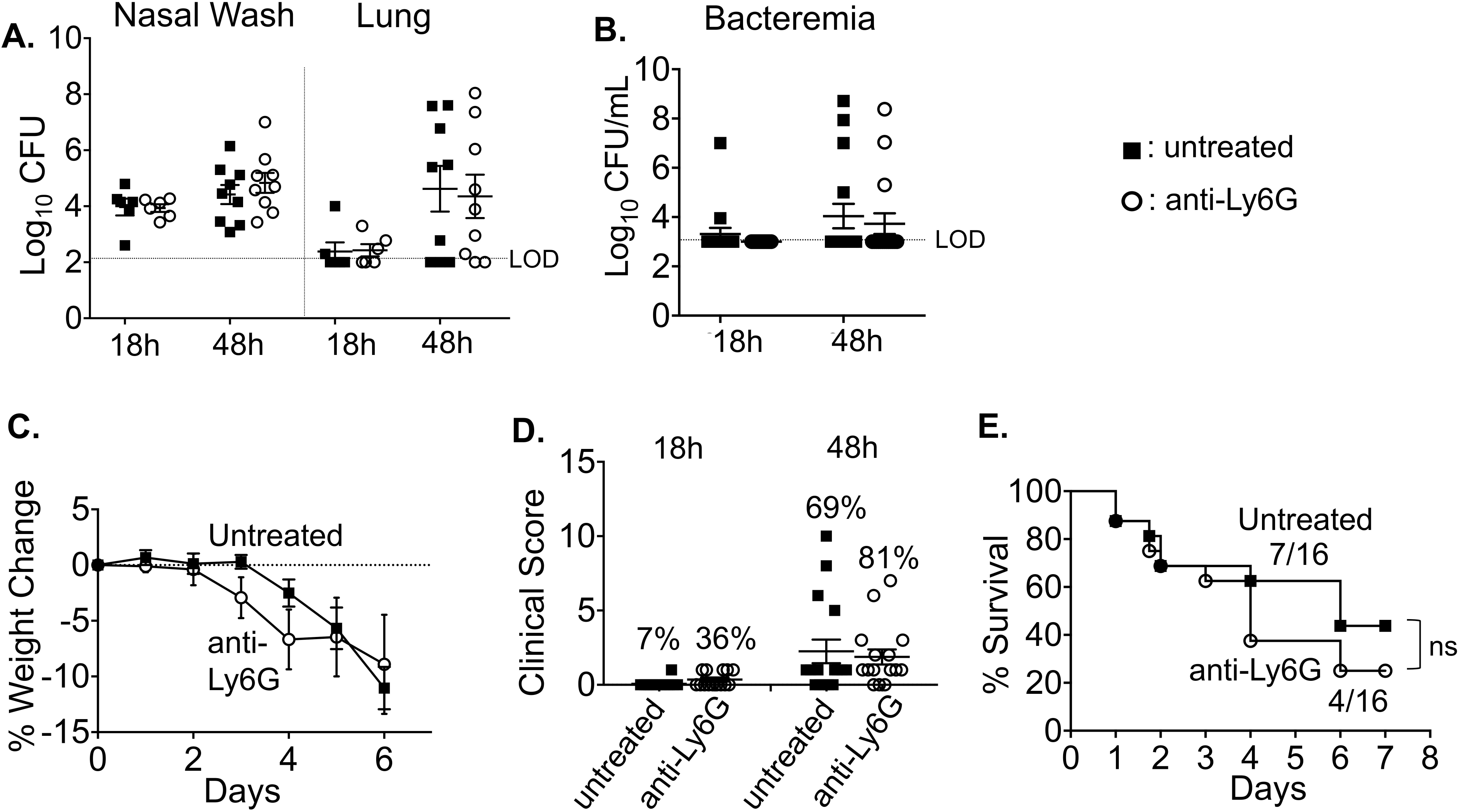
PMN depletion may have a small effect on the course of disease during IAV/*S. pneumoniae* co-infection. 8-10 weeks old C57BL/6 mice were intraperitoneally (i.p.) injected with anti-Ly6G (clone 1A8) antibodies to deplete neutrophils or mock treated. The antibodies were given daily from day −3 to day 1, and every other day from day 3 to the end of each experiment (with respect to IAV-infection). (A) At 18 and 48 hours post IAV infection, bacterial numbers in the nasal wash and lungs were determined. Pooled data from three separate experiments with a total of n=6 mice per experimental condition at 18h and n=9 mice per condition at 48h are shown. Bacteremia (B) and weight loss (C) was monitored over time. (D) Mice were blindly scored for symptoms of diseases at 18 and 48 hours post IAV-infection. The percentages indicate number of mice with clinical sickness (clinical score above 1). (E) Survival was monitored over time where the fractions denote survivors over total number of mice at 8 days post IAV infection. (C-E) Data are pooled from three separate experiments with n=16 mice per group. Statistically significant differences were determined by Student’s t-test for bacterial burden and clinical score and the log-rank (Mantel-Cox) and *ns* = not significant

### Aging increases susceptibility to IAV/*S. pneumoniae* co-infection

We next tested if the new co-infection mouse model (Fig. 1A) recapitulates the age-associated increase in susceptibility to secondary pneumococcal pneumonia. Old (20-24 months) B6 mice were inoculated i.n. with 5×10^6^ CFU of biofilm grown S. *pneumoniae* TIGR4 and 48 hours later were infected with 500 PFU IAV i.n. plus 20 PFU i.t. When compared to young co-infected controls, old mice displayed significantly more severe signs of disease, as indicated by a higher average clinical score (Fig. 3A). While seven of 16 (∼44%) young mice showed clinical symptoms (i.e., clinical score greater than 1; Fig. 3A), all 13 old mice showed at least some degree of illness by day 2 post co-infection (*p* = 0.0012, by Fisher’s exact test). Furthermore, whereas only 25% (4 out of 16) young mice had a clinical score greater than 2, which is indicative of more severe disease, 92% (12 out of 13) aged mice fell into this category (*p* = 0.005, by Fisher’s exact test). In addition, old mice died at a significantly accelerated rate. By day 2 post co-infection, 60% of old mice had succumbed to the infection compared to only 25% of young mice. Differences in survival were observed at each successive time point, and at the end of the experiment on day 8, only 14% of old mice remained alive compared to 50% of young mice (Fig. 3B). Importantly, the accelerated death observed in co-infected old mice was not observed in old mice infected with *S. pneumoniae* alone or IAV alone (Fig S4).

**Figure 3.**
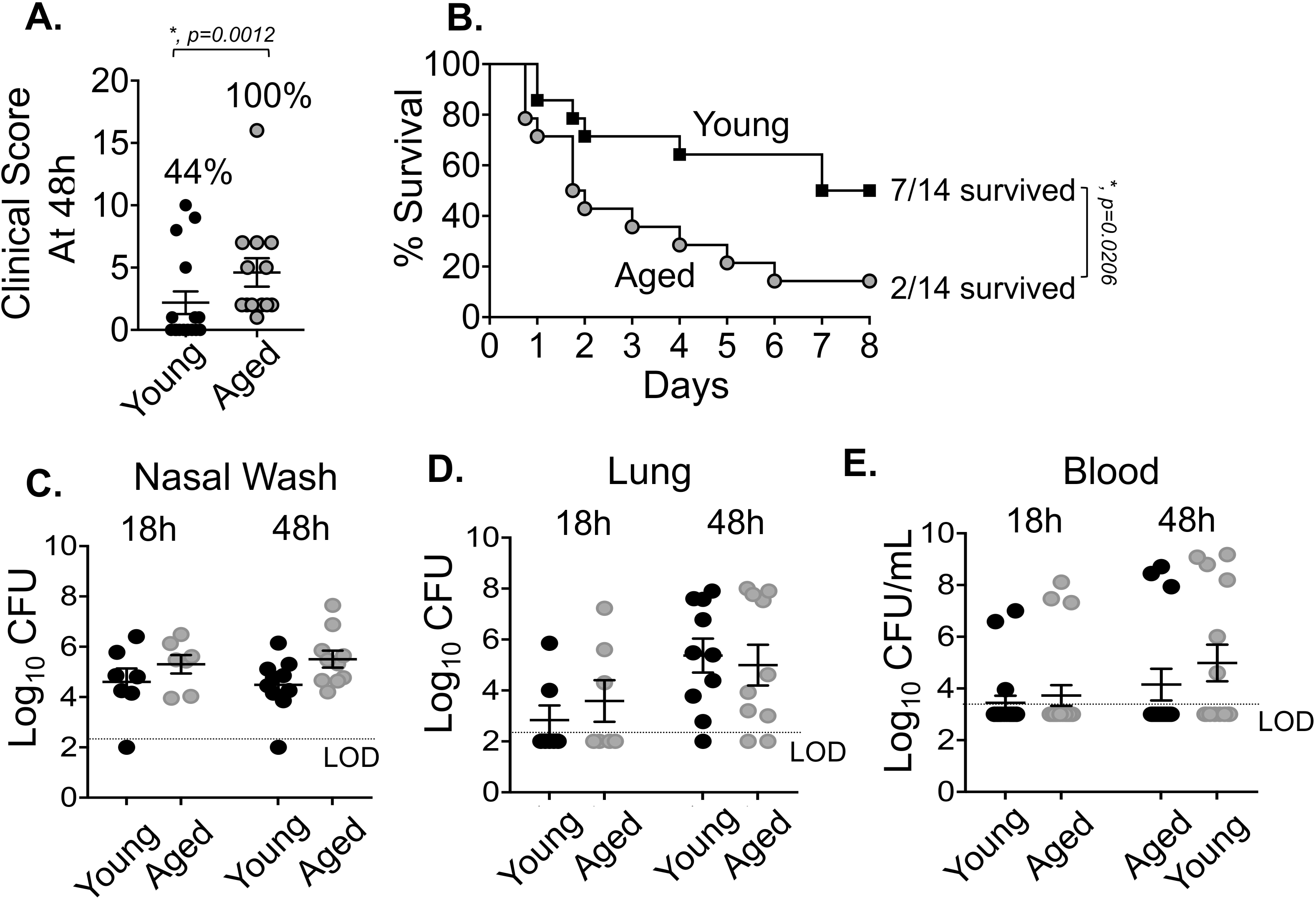
Aging increases susceptibility to IAV/*S. pneumoniae* co-infection. Young (8-10 weeks) and aged (18-24 months) C57BL/6 male mice were co-infected with *S. pneumoniae* TIGR4 i.n. and Influenza A virus PR8 i.n. and i.t. (as in Figure 1A). (A) Clinical score of co-infected young and aged mice at day 2 post IAV-infection is shown; the percentage of mice with demonstrable illness is indicated. #, indicate statistical significance by Fisher’s exact test. (B) Survival of co-infected young and aged mice were monitored over time with fractions denoting survivors over total of mice. Data are pooled from four experiments with n=14 mice per group. Asterisks indicate statistical significance by the log–rank (Mantel-Cox) test. (C-E) Bacterial burdens in the nasal wash, lungs, and blood were determined at 18 and 48 hours post IAV inoculation. The mean +/-SEM pooled from three separate experiments are shown with n=7 mice per condition at 18h and n=10 mice per condition at 48h. LOD denotes the limit of detection. For clinical score (A), data shown are pooled from all mice from experiments shown in (B) and (C-E) that had survived up to that timepoint (i.e., n= 13 for old and n= 16 for young).

We next tested whether these differences could be attributed to increased bacterial loads in the nasopharynx, lungs, or blood. As old mice got sicker at earlier time points after IAV co-infection, with the majority succumbing by day 4, we compared bacterial burden across age groups at 18 and 48 hours after IAV co-infection. We found no significant differences in the numbers of pneumococci in nasopharyngeal washes, pulmonary homogenates, or blood at either time point (Fig 3C-E). Taken together, these findings suggest that with aging there is an increased susceptibility to co-infection and an accelerated course of disease that could not be attributed to a more rapid bacterial dissemination or higher bacterial loads in the nasopharynx, lung, or bloodstream.

### Aging is associated with more rapid lung inflammation

To determine if the accelerated rate of death examined in co-infected old mice was due to more lung damage, we analyzed H&E-stained lung sections for alveolar congestion, hemorrhage, alveolar thickness, neutrophils and lymphocytic infiltration (Fig. 4A). We found that the alveolar spaces of both uninfected old and young mice were clear and free of inflammatory or red blood cells (Fig. 4A). At 18h hours post co-infection, the lungs of young mice did not show any overt signs of disease (Fig. 4A). In contrast, co-infected old mice had significant lung pathology by 18 hours post infection, including a loss of alveolar architecture, and spotty inflammation consisting of infiltrates composed of neutrophils, alveolar macrophages and mononuclear cells that were mixed with red blood cells (Fig. 4A). At 48 hours post-infection, there were clear signs of lung pathology in both young and old co-infected mice (Fig. 4A).

**Figure 4.**
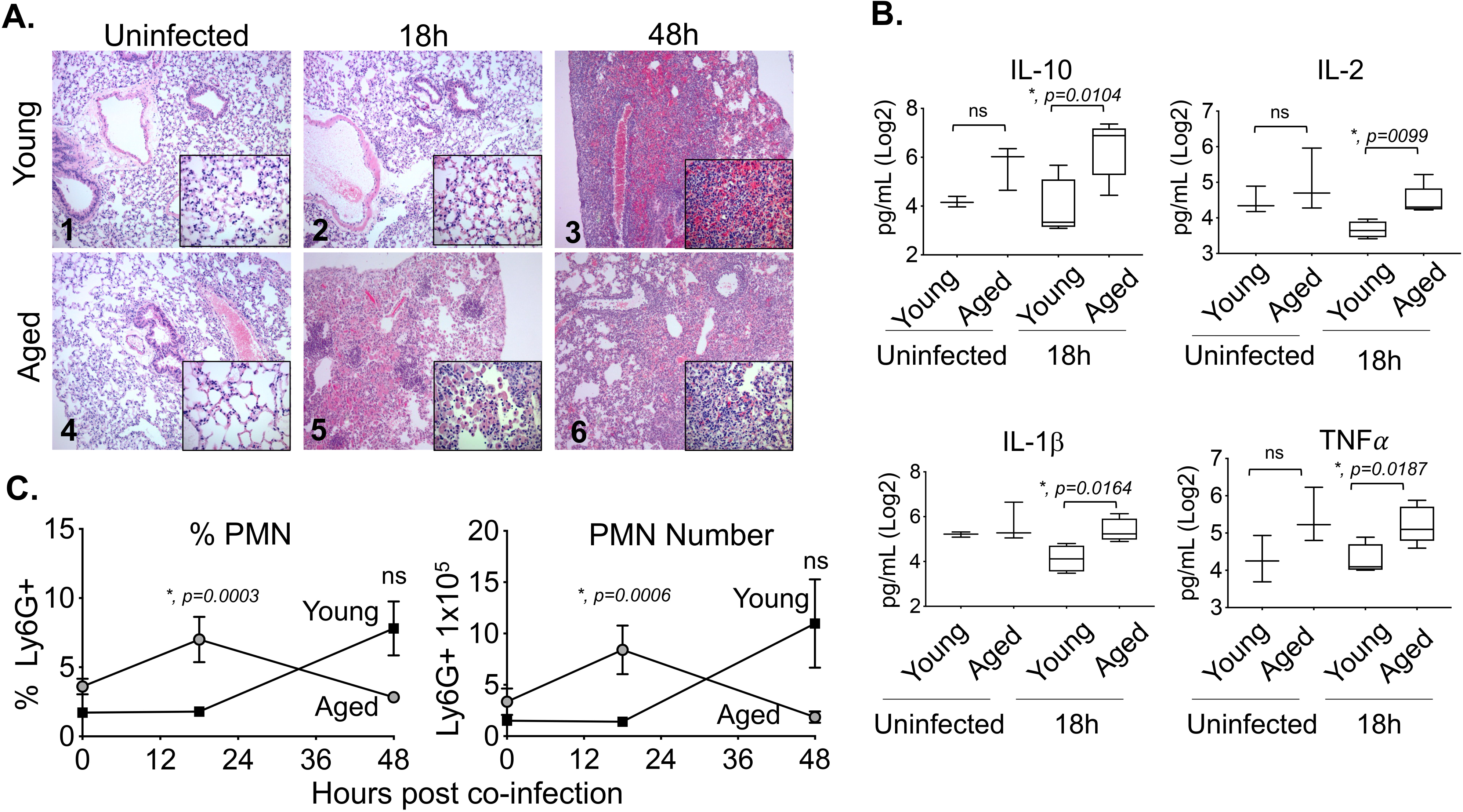
Aging is associated with more rapid lung inflammation. Young and aged C57BL/6 male mice were co-infected with S. *pneumoniae* and Influenza A virus PR8. 18 and 48 hours following IAV-infection (see experimental design in Fig. 1A), the lungs were harvested. (A) Lungs were stained with Hematoxylin and Eosin and shown are representative photographs at 100x or 400X (in inset). (B) Cytokines in the supernatants of lung homogenates of young (n=4) or aged (n=5) mice at 18 hours post co-infection were measured by multiplex ELISA. Asterisks represent statistical significance by Student’s t-test. (C) The percentages and total number of PMNs (Ly6G+) in the lungs was measured by flow cytometry. Young mice are represented by open bars and aged mice by shaded bars. The mean +/-SEM pooled from three separate experiments are shown where data are pooled from twelve mice per age group, except for 48h post-co-infection, where due to the kinetics of disease, data from only 7 surviving old mice are shown. Statistically significant differences determined by Student’s t-test are indicated by asterisks.

Next, we tested the levels of inflammatory cytokines in co-infected young vs old mice. No significant differences between age groups were detected in baseline (uninfected) levels of any of the cytokines tested between the age groups (Fig 4B and not shown). However, consistent with the enhanced PMN influx at 18 h post-infection in aged mice, old mice had significantly higher levels of IL-10, IL-2, IL-1β and TNF*α* (Fig 4B) compared with young mice. Levels of IL-12p70, IL-17, IL-6 and IFN*_γ_*, were slightly but not significantly elevated 18 hours post co-infection (Fig S5A). By 48 hours there were no significant differences between young and old mice in cytokine levels except for IFN*_γ_*, which was higher in young mice (Fig S5B).

We previously found that mice suffering from exacerbated PMN-mediated pulmonary inflammation during pneumococcal pneumonia did not display higher bacterial burdens in their lungs (27) despite a higher likelihood of severe disease (19, 20, 27). To test whether PMN influx was also higher in old co-infected mice, we measured the percentage and number of pulmonary PMNs (Ly6G^+^) by flow cytometry. We found that old mice had significantly (6-fold) higher percentages and numbers of PMNs in their lungs as compared to young controls at 18 hours post co-infection (Fig 4C). By 48 hours, most aged mice had succumbed to the infection (Fig. 3B), confounding interpretation of PMN numbers at this time point; PMN percentages and numbers appeared to be higher in young mice, but the differences were not statistically significant (Fig. 4C). Macrophages, which are important for host resistance to *S. pneumoniae/IAV* co-infection (49) and display age-driven changes (60, 61), displayed no significant age-dependent differences in either percentage or number at 18 or 48 hours after infection (Fig. S6). Taken together, these findings demonstrate that aging is associated with earlier pulmonary inflammation and damage following co-infection, which may contribute to the accelerated death observed in this mouse group.

### Aging and IAV infection diminish the ability of PMNs to kill *S. pneumoniae ex vivo*

Aged, co-infected mice experience an accelerated rate of pulmonary inflammation but bacterial loads in the lungs of aged mice were not lower than in young mice, indicating that PMN infiltration is not associated with bacterial clearance. Both aging (24) and IAV infection (62) have been reported to diminish antibacterial function of PMNs. To assess the ability of PMNs to kill *S. pneumoniae* in our co-infection model, we used a well-established opsonophagocytic (OPH) killing assay (19, 63). We first compared the bactericidal activity of bone marrow-derived PMNs from young or aged mice. The percentage of bacteria killed upon incubation with PMNs for 45 or 90 minutes was determined by comparing surviving CFU to no PMN control reactions at the same timepoint. We found that as previously reported for humans (23) and mice (24) the ability of PMNs isolated from uninfected old mice to kill pneumococci was reduced 5-fold compared to young controls regardless of the duration of infection (Fig. 5).

**Figure 5.**
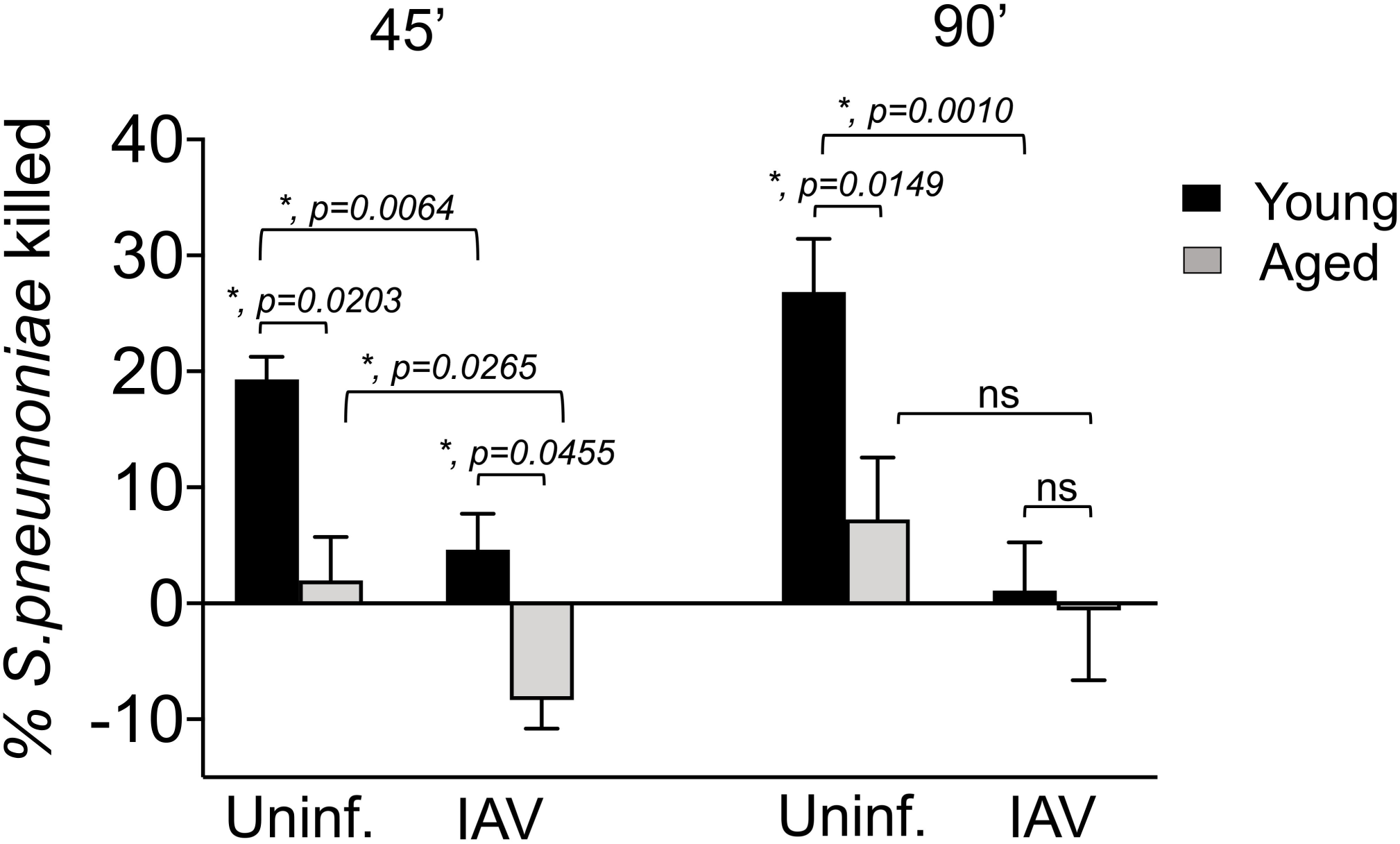
Aging and IAV infection diminish the ability of PMNs to kill *S. pneumoniae ex vivo*. PMNs were isolated from bone marrow of young (8-10 weeks) and aged (18-24 months) C57BL/6 male mice that were mock-infected (uninf.) or singly infected with IAV (i.n. + i.t.) for 2 days. PMNs were incubated with *S. pneumoniae* pre-opsonized with homologous sera from the same mouse for 45 or 90 minutes at 37°C. The percentages of *S. pneumoniae* killed upon incubation with PMNs were determined with respect to a no PMN control. Data shown represent the means +/-SEM pooled from two experiments (n=3 mice per group per timepoint) where each condition was tested in quadruplicates per experiment. Asterisks represent statistical significance as determined by Student’s t-test.

We next examined the bactericidal activity of PMNs isolated from young or old mice 2-days after i.t./i.n. IAV inoculation. As previously reported (47–50), PMNs from young IAV infected mice had a slight (2-fold) but significant reduction in their ability to kill pneumococci as compared to PMNs from uninfected controls (Fig. 5). Strikingly, IAV infection diminished the ability of PMNs from old mice to kill *S. pneumoniae*; instead, PMNs from IAV infected old mice promoted a slight increase in bacterial numbers (Fig. 5). These findings suggest that IAV infection completely abrogates the ability of PMNs from old mice to kill *S. pneumoniae*.

### Aging and prior colonization with *S. pneumoniae* result in impaired IFN-**α** production and higher viral burden in the lungs

Finally, we investigated whether aging and/or bacterial co-infection had an impact on antiviral responses. Old or young B6 mice were inoculated i.n. with 5×10^6^ CFU of biofilm grown S. *pneumoniae* TIGR4 or mock challenged with PBS and 48 hours later were infected with 500 PFU IAV i.n. and 20 PFU i.t. At 48 hours following viral infection, we compared viral burden in the lung across age groups. We found that bacterial colonization resulted in 10-fold (and statistically significant) higher pulmonary viral loads when compared to mock-colonized controls, regardless of host age (Fig. 6A). Further, we found that in co-infected hosts, aging was associated with significantly increased viral loads in the lungs (Fig. 6A), suggesting that enhanced viral loads and impaired antiviral defenses contribute to the differences in clinical manifestation across host age.

**Figure 6.**
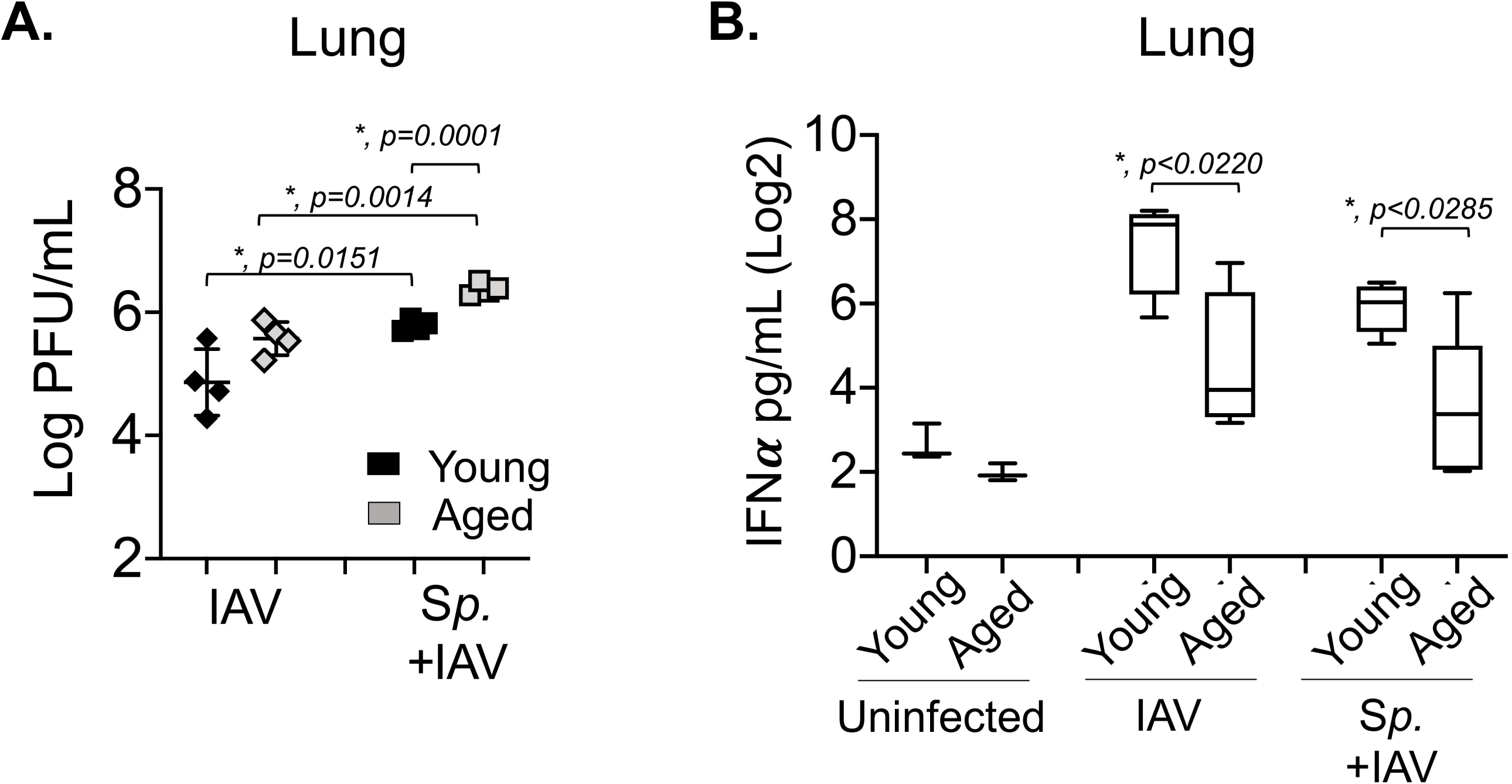
Aging and prior colonization with *S. pneumoniae* result in impaired IFN-α production and higher viral burden in the lungs. Young and aged C57BL/6 male mice were either co-infected with S. *pneumoniae* and Influenza A virus PR8 (*Sp* + IAV), challenged with virus alone (IAV) or mock challenged with PBS (uninfected) (as in Figure 1A). Two days post IAV-infection, (A) viral burdens in the lungs were determined and (B) levels of IFN-α in the supernatants of lung homogenates were measured by ELISA. Data from n=4 mice per group are shown. Statistically significant differences were determined by Student’s t-test.

To test whether bacterial colonization and aging impair antiviral immune responses, we measured the levels of IFN-α, a cytokine crucial for antiviral defense (64). We found that in young mice, despite the higher viral burden (Fig. 6A), prior bacterial colonization resulted in a 3-fold (but not statistically significant) decrease in IFN-α production in the lungs in response to IAV challenge (Fig. 6B). Notably, aging was associated with (statistically significant) 5- and 3-fold lower levels of IFN-α correspondingly in mice infected with IAV alone or co-infected with IAV and *S. pneumoniae.* No differences in IFN-α production were observed in the sera (Fig S7). These findings suggest that both pneumococcal infection and aging blunt antiviral responses in this mouse model.

## DISCUSSION

*S. pneumoniae* remains a leading cause of secondary bacterial pneumonia following influenza A virus infection and is associated with severe disease (30, 34–36), particularly in the elderly (52). The majority of *S. pneumoniae/*IAV co-infection experimental studies have delivered bacteria into the lungs of mice pre-infected with IAV to reveal changes in the host lungs and immune response that are crucial for priming invasive pneumococcal disease (34, 41, 44–50). In this study, to investigate the transition of *S. pneumoniae* from colonizer to pathogen upon IAV co-infection, a process that has just started to be elucidated (4, 37), we have developed a modified murine infection model that recapitulates this transition and results in severe clinical disease. In a previously established model, female BALB/cByJ mice were first colonized intra-nasally with biofilm-grown pneumococci and then infected with IAV by delivering the virus to the nasopharynx (4, 40). This model showed that changes in the host environment in response to viral infection triggers the dispersal of pneumococci from colonizing biofilms and their spread to the lower respiratory tract (4). Importantly, the dispersed bacteria expressed higher levels of virulence factors required for infection, thus, rendering them more highly pathogenic (4). Nevertheless, upon dispersion following *in vivo* IAV infection, although bacterial migration to the lung was detected, the burden was less than 100 CFU per lung and the mice did not suffer overt disease. When we used this model to co-infect male C57BL/6 (B6) mice, we also observed a significant increase in dispersed bacteria in the nasopharynx, but no disease and only a transient presence of *S. pneumoniae* in the lungs of B6 mice. Here we performed experiments in male instead of female mice due to both the easier availability of aged male animals and the documented higher rate of pneumococcal pneumonia in men compared to women (65, 66).

Previous work of pneumococcal inoculation of IAV-infected lungs showed viral infection to be crucial for creating an environment in the lungs that is more permissive for bacterial infection (41, 44–50). The relatively low bacterial burden in co-infected male B6 mice observed here upon IAV inoculation only by the i.n. route suggested that the lung environment was not sustaining the bacteria. It is possible that human IAV/*S. pneumoniae* co-infection involves viral infection of not only the upper respiratory tract, but the lower respiratory tract as well (54, 55). Therefore, we co-infected *S. pneumoniae* colonized B6 mice with IAV by delivering the virus not only i.n. to infect the nasopharynx, but also i.t. to ensure infection of the lungs. This modified model recapitulated the increase in non-adherent pneumococci in the nasopharynx observed upon viral co-infection (4, 67, 68), and resulted in bacterial spread into the lungs and circulation that increased over time. Importantly, this mode of dual infection recapitulated both the increased colonization burden (69, 70) as well as the clinical signs of severe disease observed in humans (30, 34–36), and resulted in death of approximately half of co-infected young controls.

Although the elderly are at higher risk for secondary pneumococcal pneumonia following IAV infection, animal studies exploring this age-driven susceptibility to co-infection are few (51, 71, 72). Here, using the modified model, we found that old mice were significantly more susceptible to *S. pneumoniae/*IAV co-infection. Old mice displayed more severe signs of disease as compared to young controls and the majority (>85%) failed to survive the co-infection. This increased susceptibility in old mice was not linked to higher bacterial dissemination from the nasopharynx, greater establishment of infection in the lungs, or systemic spread into the circulation at the time points tested. Susceptibility to viral infection alone, as measured by weight loss within the first 5 days following IAV, was also similar between the age groups. Rather, the age-driven susceptibility to co-infection was associated with earlier and more severe pulmonary inflammation. Production of pulmonary cytokines is elevated in young mice infected with *S. pneumoniae* at 7 days after IAV inoculation (73). Further, old mice display changes in the expression of pattern recognition receptors in the lungs leading to altered inflammatory responses (51). Similar to previous studies (51), we found here that co-infected old mice had higher levels of TNF*α* compared to young controls. However, in contrast to previous reports that found reduced NLRP3 inflammasome expression in the lungs and lower production of IL-1β, we found higher levels of IL-1β in old vs. young mice. This may be accounted for by differences in expression of bacterial factors. The expression of pneumolysin, which was found to activate NLRP3 inflammasomes and lead to production of IL-1β (74), was elevated in pneumococci dispersed from biofilms upon IAV infection (75) and therefore may have primed the IL-1β production we observed in old mice.

We previously found that PMNs were key determinants of disease during primary pneumococcal pneumonia and are required to initially control bacterial numbers (20, 76). Therefore, we explored here the role of PMNs in *S. pneumoniae/*IAV co-infection. Similar to other studies, we found that PMNs are recruited to the lungs of co-infected young mice (49). Previous reports indicate that the PMN-mediated anti-pneumococcal function in IAV-infected mice (62) and humans (70) is progressively reduced over time. For example, in mice, PMNs demonstrably contribute to host defense at 3 days but not at 6 days post-infection (62). Nevertheless, PMN depletion showed that these cells are important for control of bacterial transmission (77) and control of pulmonary bacterial numbers (49) in IAV co-infected mice. Similarly, here we found that PMN depletion starting prior to bacterial colonization and continuing throughout viral co-infection appeared to slightly worsen disease progression, with slightly greater average weight loss at days 3 and 4 after viral inoculation and a greater proportion of PMN-depleted mice succumbing to co-infection. However, PMN depletion had no significant effect on the number of dispersed *S. pneumoniae* in the nasopharynx or their spread to the lungs and blood. It is possible that, in this model, IAV may rapidly impair PMN function, limiting their efficacy even in PMN-replete mice. In fact, we found that within 2 days following IAV infection, the ability of bone marrow-derived PMNs to kill *S. pneumoniae* was significantly blunted in young mice. Alternatively, the apparent inability of PMNs to limit bacterial numbers in this model could be due to the enhanced virulence of pneumococci dispersed from the nasopharyngeal environment (4) compared to broth-grown bacteria typically used in other models (62). In this study we found that aging was associated with earlier influx of PMNs into the lungs of co-infected mice. We previously demonstrated that excessive PMN influx into the lungs is detrimental for the ability of old mice to control invasive disease following primary pneumococcal pneumonia (19) and that PMN depletion 18 hours after infection boosted host survival (20). Similarly, greater PMN influx into the lungs of old mice singly infected with IAV was associated with host mortality, and depletion of these cells six days following viral infection significantly boosted host survival (78). Uncontrolled PMN influx can result in tissue damage, disruption of gaseous exchange and pulmonary failure. In fact, it was reported that in IAV singly infected old mice, PMNs enhanced lung inflammation and damage and their depletion reduced the levels of inflammatory IL-1β and TNFα(78). Therefore, the early increase in these inflammatory cytokines and lung damage we observed here in *S. pneumoniae/*IAV co-infected old mice may be driven by the elevated levels of pulmonary PMNs.

The age-associated increased levels of pulmonary IL-10 during *S. pneumoniae/*IAV confection observed here may also contribute to the enhanced susceptibility of old mice to co-infection. IL-10 levels are elevated in co-infected young mice compared to those singly infected with *S. pneumoniae* (79). We showed that blocking this cytokine boosts PMN anti-bacterial function by inhibiting ROS production (76), and van der Poll and coworkers demonstrated that inhibition of IL-10 restores resistance of vulnerable hosts to primary pneumococcal pneumonia (79).

The more severe disease observed in old mice also correlated with higher viral lung burden during both co-infection with *S. pneumoniae* and IAV and single infection with IAV, although the difference reached statistical significance only in the former. Production of type I IFN, which is crucial for control of viral replication, is dysregulated during aging (80) and we found that pulmonary IFN-α levels of co-infected or singly infected old mice were significantly lower compared their young counterparts. Hence, in this model, enhanced disease associated with aging may be a reflection of a combination of an exuberant PMN response and a muted type I interferon response.

While many studies have examined the effect of IAV on secondary pneumococcal pneumonia, the effect of bacterial colonization on viral infection has been less explored. We found that, regardless of host age, prior colonization with pneumococci resulted in significantly higher viral pulmonary loads. In experiments involving two different hosts (ferrets or cotton rats) and two different viruses (IAV and Human RSV, respectively), nasopharyngeal colonization of donors with *S. pneumo*niae promoted viral transmission (81, 82). In the latter (rat-RSV) model, prior pneumococcal colonization enhanced viral infection of the upper respiratory tract but did not promote infection of the lungs (82). A human experimental pneumococcal colonization study showed that prior colonization with pneumococci did not increase the burden of a live attenuated influenza virus, but reduced pro-inflammatory immune responses in the nasopharynx and significantly blunted antiviral IgG production in the lungs (83). Hence, prior colonization with pneumococci may impair antiviral defenses. Indeed, we found that in spite of their approximately one log higher viral lung burden compared to singly infected mice, *S. pneumo*niae-pre-colonized young and old mice respectively displayed 5 and 3-fold lower levels of pulmonary IFN-α compared to non-colonized controls.

In summary, here we modified existing murine models to establish an experimental system that reflects the transition of *S. pneumoniae* from asymptomatic colonizer to invasive pulmonary pathogen upon IAV co-infection. In this model, IAV triggers the transition of a pathobiont from a commensal to a pathogenic state through modification of bacterial behavior in the nasopharynx, enhancement of bacterial colonization in the lung, and compromise of PMN-mediated anti-bacterial immunity. In turn, pneumococci modulate antiviral immune responses and promote IAV infection of the lower respiratory tract.

Importantly, this model recapitulates the susceptibility of aging to co-infections. Moving forward, this multi-step model can be used to dissect both the multiple phases of pneumococcal disease progression from commensals to pathogens and the complexity of viral/bacterial interactions within different hosts, thus helping inform specialized treatment options (67) tailored to the susceptible elderly population.

## MATERIALS AND METHODS

### Mice

Young (8-10 weeks) and aged (18-24 months) male C57BL/6 mice were purchased from Jackson Laboratories (Bar Harbor, ME) and the National Institute of Aging. Mice were housed in a pathogen-free facility at Tufts University. All procedures were performed in accordance with Institutional Animal Care and Use Committee guidelines. The number of mice included in each experiment is based on power analysis. At least 6 mice per group for determination of bacterial burden and 12 mice per group for monitoring survival were included to sufficiently power our studies. In each experiment, an equal number of mice per group were planned. However, slight variations in the number of mice per group at later time points of several experiments occurred due to the required euthanasia of several mice in highly susceptible experimental groups.

### Bacterial biofilms

NCI-H292 mucoepidermoid carcinoma cells (H292) were grown in 24-well plates in RPMI 1640 media with 10% FBS and 2mM L-glutamine until confluent. Cells were washed with 1x PBS and fixed in 4% paraformaldehyde for 1 hour on ice. *S. pneumoniae* TIGR4 (kind gift from Andrew Camilli) were grown on Tryptic Soy Agar plates supplemented with 5% sheep blood agar (blood agar plates) overnight, then diluted and grown in chemically defined liquid medium (CDM) (4, 40) supplemented with Oxyrase until OD_600nm_ of 0.2. Bacteria were diluted 1:1000 in CDM and seeded on the fixed H292 cells. The, bacteria/H292 cells were incubated at 34°C/ 5% CO_2_ and media was changed every 12 hours. At 48 hours post-infection, the supernatant containing planktonic bacteria (non-adherent to NCI-H292 cells) cells was discarded, the cells gently washed with PBS and adherent biofilms collected in fresh CDM by vigorous pipetting. Biofilm aliquots were then frozen at −80°C in the CDM with 25% (v/v) glycerol.

### Intranasal inoculation

Before use, biofilm aliquots were thawed on ice, washed once and diluted in PBS to the required concentration. The mice were restrained without anesthesia and infected i.n. with 10μl (5×10^6^ CFU) of biofilm grown *S. pneumoniae*. The inoculum was equally distributed between the nostrils with a pipette. This method of inoculation results in pathogen delivery limited to the nasal cavity, without accessing the lower tract in C57BL/6 mice (58). Bacterial titers were confirmed by serial dilution and plating on blood agar plates. To ensure stable colonization of the biofilm in the nasopharynx, groups of mice were euthanized at 18 and 48 hours post inoculation, and the nasal washes and tissue were collected and plated on blood agar plate for enumeration of *S. pneumoniae*.

### Viral infection

The mouse-adapted H1N1 Influenza A virus PR8 (A/PR/8/34) was - obtained from Dr. Bruce Davidson (4, 40) and stored at −80 ☐C. Before use, viral aliquots were thawed on ice, diluted in PBS and used to inoculate mice. At 48 hours following bacterial inoculation, mice were infected i.n. with 10μl of 500 plaque forming units (PFU) of virus by pipetting the inoculum into the nostrils of non-anesthetized mice. Following i.n. inoculation, mice were lightly anesthetized with isoflurane and challenged i.t. with 20 PFU of virus in a 50μl volume pipetted directly into the trachea with the tongue pulled out to facilitate delivery (20).

### Clinical scoring and bacterial burden

Following co-infection, mice were monitored daily and blindly scored for signs of sickness including weight loss, activity, posture and breathing. Based on these criteria, the mice were given a clinical score of healthy [0] to moribund [25] modified from what was previously described (59). Any mice displaying a score above 9 are humanely euthanized in accordance with our protocol. Mice were euthanized at indicated time points and the lung, nasal lavages and blood collected and plated on blood agar for enumeration of bacterial loads, as previously described (20). For collection of sera, blood was collected via cardiac puncture into Microtainer^®^ tubes (BD Biosciences) and centrifuged at 7607xg for 2 minutes to collect serum as per manufacturer’s instructions.

### Depletion of PMNs

Mice were intraperitoneally (i.p.) injected with 100 lμ (50μg/mouse) of anti-Ly6G clone 1A8 (BD Biosciences) to deplete neutrophils. Mice were injected daily with the depleting antibodies starting one day prior and ending two days after bacterial inoculation, followed by every other day from day 1 post viral co-infection to the end of each experiment. Treatment resulted in >90% neutrophil depletion as described below.

### Cell isolation and Flow Cytometry

Mice lungs were harvested, washed in PBS, and minced into small pieces. The sample was then digested for 45 minutes with RPMI 1640 1X supplemented with 10% FBS, 1 mg/ml Type II collagenase (Worthington, Lakewood, NJ), and 50 U/ml deoxyribonuclease I to obtain a single-cell suspension as previously described (20). The red blood cells were lysed using ACK lysis buffer (Gibco). Cells were then stained with anti-mouse Ly6G clone 1A8 (BD Biosciences), F480 clone BM8 (BioLegend), CD11c clone N418 (eBioscience), and CD11b clone M1/70 (eBioscience). For neutrophil depletion, cells isolated from the lungs at 18 and 48 h post co-infection were also stained with either Ly6G clone 1A8 or RB6 clone RB6-8C5 (BioLegend) antibodies to confirm cell depletion. The fluorescence intensities were measured on BD LSR II Flow Cytometer at Tufts FACS Core Facility (Boston, MA) to capture at least 25,000 cells and analyzed using FlowJo.

### Isolation of PMNs and Opsonophagocytic Killing Assay (OPH)

Femurs and tibias of uninfected mice were collected and flushed with RPMI, supplemented with 10% FBS and 2 mM EDTA to obtain bone marrow cells. Neutrophils were isolated by density gradient centrifugation, using Histopaque 1119 and Histopaque 1077 as previously described (84). The neutrophils were resuspended in Hank’s (Gibco) buffer/0.1% gelatin with no Ca+ or Mg+ and tested for purity by flow cytometry using anti-Ly6G antibodies (eBioscience) where 85-90% of enriched cells were Ly6G+. The ability of neutrophils to kill bacteria was measured using a well-established opsonophagocytic (OPH) killing assay as previously described (20). Briefly, 2.5×10^5^ neutrophils were incubated in Hank’s buffer/0.1% gelatin with 10^3^ CFU of *S. pneumoniae* pre-opsonized with 3% mouse sera. The reactions were incubated in flat bottom 96-well non-binding plates for 45 minutes at 37°C. Each group was plated on blood agar to enumerate viable CFU. Percent bacterial killing was calculated in comparison to a no PMN control under the same conditions.

### Histology

Whole lungs were harvested from groups of mice at 18 hours and at 48 hours post co-infection and fixed in 10% neutral buffered formalin for 2 days. The tissues were then embedded in paraffin, sectioned at 5 μm and stained with Hematoxylin and Eosin at the Animal Histology Core at Tufts University. Sections of lung from three mice per group were imaged using a Nikon Eclipse E400 microscope. Photomicrographs were captured using a SPOT Idea 5.0-megapixel color digital camera and SPOT software. Histopathologic scoring was performed by a board-certified anatomic pathologist experienced in murine pathology, from 0 (no damage) to 4+ (maximal damage) for alveolar congestion, hemorrhage, alveolar thickness, neutrophils, and lymphocytic infiltration (85).

### Cytokine Analysis

Frozen lung homogenates and serum samples were thawed on ice and mixed by gentle vortexing. Cytokines in the lungs and serum samples were measured using Mouse Cytokine 8-Plex Array (Quanterix, Billerica, MA) following the manufacturer’s instructions. Levels in lung supernatants and serum samples were measured using the Cirascan at the Imager at Tufts University Genomic Core (Boston, MA) and analyzed by the Cirascan/Cirasoft program. Qlucore Omic Explorer (version 3.5) was used for the generation of lung cytokine box plots. Concentrations of cytokines (IL-10, IL-2, IL-1β TNF*α*, IL-6, IFN*γ*, IL-17 and IL-12p70) were log-transformed, and displayed as Log_2_ pg/ml. IFNα was measured using Mouse IFN-alpha ELISA kit (R&D system, MN). following the manufacturer’s instructions.

### Plaque assay

Madin-Darby Canine Kidney (MDCK) cells were grown overnight in 12 well plates at 2×10^5^cells/well in DMEM media +10% FBS. Cells were washed twice with 1xPBS and incubated with serial dilutions of viral inoculum or lung homogenates in DMEM supplemented with 0.5% low endotoxin BSA (Sigma-Aldrich) for 50 minutes in 37 ☐C 5% CO2 incubator. The plates were shaken every 10 minutes during the incubation and then washed twice with 1xPBS. 2mL of 2.4% of avicel overlay (FMC) was then added onto the infected cells and were incubated for 3 days in 37 ☐C with 5% CO2 incubator. After 3 days, avicel overlays were removed and cells were washed with 1x PBS and fixed in 4% paraformaldehyde for 30 minutes at room temperature. 1% crystal violet were then added for 5 minutes to count plaques.

### Statistical Analysis

Statistical analysis was performed using Graph Pad Prism7. CFU data were log-transformed to normalize distribution. Data are presented as mean values +/-SEM. Significant differences (*p* < 0.05) were determined by Student’s t-test. Differences in fractions of mice that got sick were measured using Fisher’s exact test. Survival analysis, including the kinetics by which mice succumb to infection, was performed using the log rank (Mantel-Cox) test. Asterisks indicate significant differences and *p* values are noted in the figures.

## ACKNOWLEDGEMENTS

We would like to acknowledge James Nicholas Lee, Summer Schmaling, and Ognjen Sekulovic for technical assistance with clinical score, virus preparation and nasal lavage respectively. We would also like to thank Andrew Camilli for bacterial strains and Marta Gaglia for protocols. They, along with Tim van Opijnen, Bharathi Sundaresh and Marcia Osburne provided important feedback on the manuscript.

## FUNDING

Research reported in this publication was supported by the National Institute On Aging of the National Institutes of Health under Award Number R21 AG064215 to E.B.G. and F31 AI122615-01A1 to S.R.; King Abdullah Scholarship Program (KASP) implemented by the Ministry of Higher Education (MOHE) under Award Number 7896504 to B.H.J.

## Notes

### Competing Interest Statement

The authors have declared no competing interest.

