## Supplemental Material for "A Murine Model for Enhancement of *Streptococcus pneumoniae* Pathogenicity Upon Viral Infection and Advanced Age"

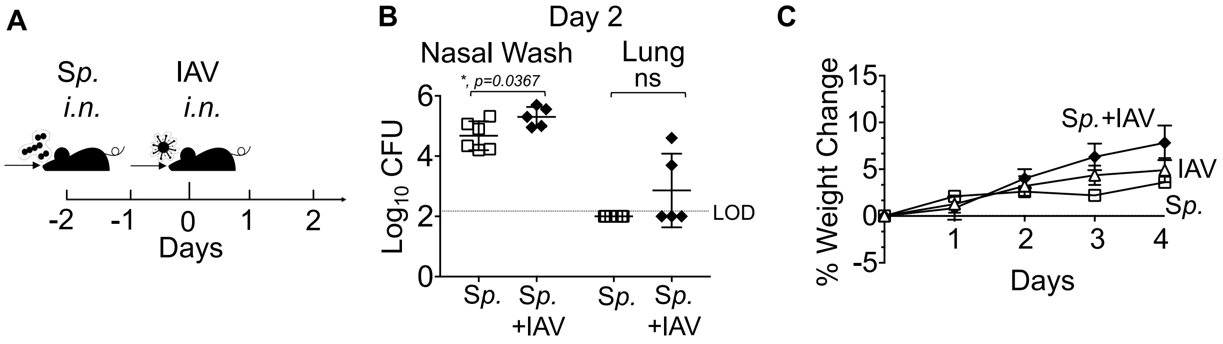


**Figure S1: Course of disease following i.n. IAV inoculation**. (A) 8-10 weeks old C57BL/6 mice were pre-colonized i.n. with biofilm generated S. *pneumoniae* TIGR4 (*Sp*) at 5x$10^6$ CFU, followed by i.n. co-infection with 500 PFU of Influenza A virus (PR8). (B) Two days post IAV-infection, mice were sacrificed and bacterial burdens in nasal lavage (left) and lung homogenates (right) were determined. Data were pooled from two separate experiments with n=6 *Sp* infected mice and n=5 co-infected mice per group. (C) Mice were also monitored daily for weight change. Data were pooled from two separate experiments with n=9 mice per group. Dotted lines represent the limit of detection. Asterisks (*) represent statistical significance, and *ns* = not significant, determined using Student’s t-test.

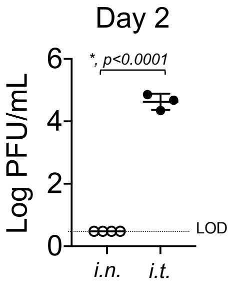


**Figure S2. Intratracheal but not intranasal delivery of IAV results in viral infection of the lungs**. 8-10 weeks old C57BL/6 mice were infected i.n. with 500 PFU of Influenza A virus (PR8) or challenged i.t. with 20 PFU of PR8. Two days later, mice were sacrificed and viral burdens in lung homogenates were determined. Data from three mice per group are shown. Dotted lines represent the limit of detection. Asterisks (*) represent statistical significance, determined using Student's t-test.

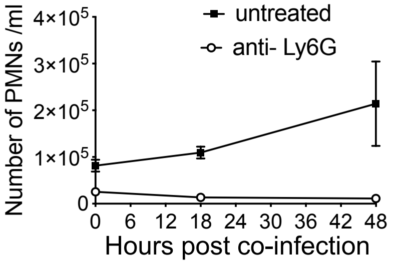


**Figure S3. PMN numbers following depletion in co-infected mice.** 8-10 weeks old C57BL/6 mice were intraperitoneally (i.p.) injected with anti-Ly6G (clone 1A8) antibodies to deplete neutrophils or mock treated. The antibodies were given daily from day -3 to day 1, and every other day from day 3 to the end of each experiment (with respect to IAV-infection). At 0, 18 and 48 hours post IAV infection, blood was collected and the number of circulating PMNs ($RB6^{hi}$) was determined by flow cytometry. Data shown represent the means +/- SEM and are representative from one of three separate experiments with n=3 mice per group.

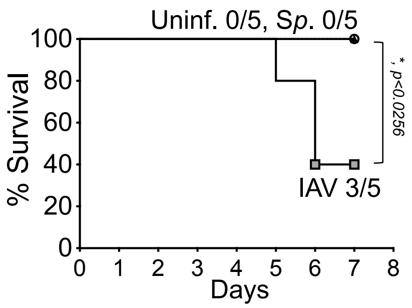


**Figure S4. Course of survival following intranasal *S. pneumoniae* inoculation or intranasal/intratracheal IAV inoculation of aged mice.** Aged (18-24 months) C57BL/6 male mice were singly infected with *S. pneumoniae* TIGR4 i.n. (*Sp*) or Influenza A virus PR8 i.n and i.t. (IAV) or mock-treated (uninfected) and survival was assessed over time and compared to uninfected controls. Fractions denote survivors over total number of mice.

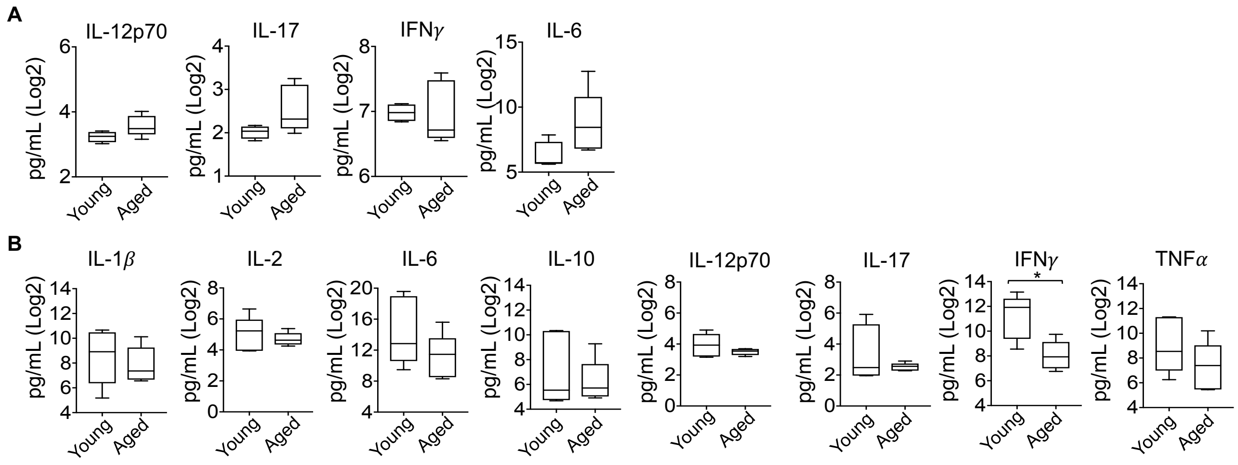


**Figure S5: IAV/*S. pneumoniae* co-infection results in no significant age-dependent differences in TH1/TH17 cytokines.** Young and aged C57BL/6 male mice were co-infected with S. *pneumoniae* and Influenza A virus PR8. Cytokines in the supernatants of lung homogenates of young (n=4) or aged (n=5) mice at (A) 18 hours and (B) at 48 hours following co-infection were measured. Asterisks (*) represent statistical significance as determined by Student's t-test.

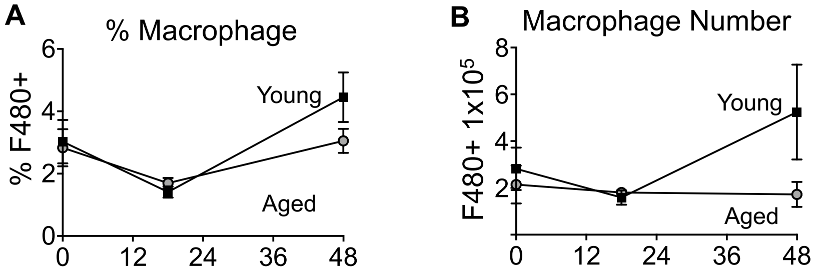


**Figure S6: IAV/*S. pneumoniae* co-infection results in no significant age-dependent differences in pulmonary macrophage influx.** The percentage (A) and total number (B) of pulmonary macrophages of co-infected young or aged mice were identified by flow cytometry using the F480 marker at the indicated times. The mean +/- SEM pooled from three separate experiments are shown where data are pooled from twelve mice per age group, except for 48 hours post-co-infection, where due to the kinetics of disease data from only 7 surviving old mice are shown.  No significant differences were detected using Student's t-test.

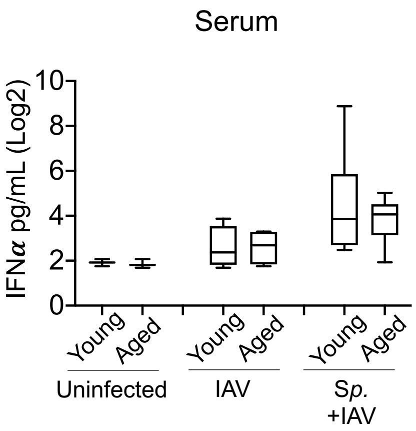


**Figure S7. IFN-α levels in the sera following infection.** Young and aged C57BL/6 male mice were either co-infected with S. *pneumoniae* and Influenza A virus PR8, challenged with virus alone or mock challenged with PBS. 48 hours following IAV-infection (see experimental design in Fig. 1A), blood was collected and serum levels of IFN-α were measured by ELISA. Pooled data from two separate experiments with n=4 mice per group are shown.
